# Virus-host interaction: investigating novel transcription factors involved in coupling HPV life cycle and epithelial differentiation

**DOI:** 10.1101/2024.06.21.599992

**Authors:** Aline Lopes Ribeiro, Valéria Talpe Nunes, Amanda Schiersner Caodaglio, Rafaella Almeida Lima Nunes, João Simão Pereira Sobrinho, Laura Sichero

## Abstract

High-risk human papillomavirus (HPV) causes almost all cervical cancer, and HPV-18 is the second most prevalent type. The HPV life cycle is intimately associated with the epithelial differentiation program, and the repertoire of cellular transcription factors (TFs) has a crucial role in coordinating viral gene expression across epithelial layers. We aimed to identify host TFs involved in the virus differentiation-dependent life cycle. We initially compared the DNA binding activity of 345 TFs in nuclear extracts of undifferentiated and high calcium-induced differentiated HaCaT cells. Next, we searched *in silico* for putative binding sites within the HPV-18 long control region (LCR) for the TFs affected by differentiation. Chromatin immunoprecipitation assays (ChIP) were used to assess direct binding of selected TFs, protein levels were evaluated by immunohistochemistry in rafts obtained from human foreskin keratinocytes (HFK) and HPV-18 E6/E7 immortalized HFK, and the impact upon LCR activity was measured in reporter assays. The binding activity of 40 TFs was influenced by differentiation among which 16 putative binding sites within the LCR were identified. We next focused our analysis in PAX6, HMGB1, FOXI1, and NFE2, and all bound to the LCR except FOXI1. Whether FOXI1 activates HPV-18 early promoter activity, PAX6, HMGB1, and NFE2 inhibit viral transcription in undifferentiated normal HFK in which these proteins are mostly expressed. Finally, HPV-18 E6/E7 impaired the normal expression of these TFs throughout epithelial layers. In conclusion, we describe the involvement of PAX6, HMGB1, NFE2, and FOXIl in coupling HPV-18 transcriptional regulation with the epithelia differentiation program, a key event for the productive HPV life cycle. Furthermore, these TFs may also be relevant for HPV-driven carcinogenesis. The identification of proteins involved in the differentiation-dependent HPV life cycle and viral induced transformation may contribute to unveiling novel therapeutic targets for HPV-related malignancies.

**Author Summary:** HPV-16 and HPV-18 respond together for over 70% of cervical cancers. HPV infection is established in basal layers cells of the stratified epithelia and the viral life cycle is highly dependent on the epithelial differentiation program. In this study, we focused on identifying host transcription factors (TFs) involved in the finely tuned regulation of the HPV-18 differentiation-dependent life cycle. We initially screened 345 cellular TFs and found 40 with differential binding activity after differentiation, among which we chose to focus on PAX6, HMGB1, FOXI1, and NFE2. Besides investigating TFs binding to HPV-18 LCR *in silico* and with immunoprecipitation assays, we evaluated protein levels throughout epithelial layers and demonstrated that the impact of these TFs upon HPV-18 early promoter activity depends on the differentiation status of cells. Finally, we show that HPV-18 oncoproteins lead to abnormal expression of these TFs. Our results contribute to a clearer understanding of pivotal mechanisms governing spatiotemporal regulation of viral early transcription and open avenues for innovative therapeutic strategies for HPV-associated diseases.

## Introduction

Human papillomaviruses (HPVs) are one of the most common sexually transmissible infections and the leading cause of virus-induced cancers worldwide [1]. HPV infections are associated with the development of almost 5% of all new cancer cases, including cancer of the uterine cervix and other anogenital sites, in addition to the oropharynx [2]. The most clinically relevant types are HPV-16 and HPV-18, responsible for more than 70% of cervical cancers [3].

HPV productive infections occur in the mucosal and cutaneous stratified squamous epithelium. The viral life cycle is highly dependent on the epithelial differentiation program [4,5] and initiates in undifferentiated dividing basal cells where HPV genomes are maintained extrachromosomally in a moderate copy number (50-100 copies per cell). As cells divide and migrate towards the epithelial surface, viral gene transcription initiates from the early promoter, termed P105 in HPV-18, enabling the expression of E6 and E7 transcripts [6]. This step is essential for E6 and E7 oncoproteins delay the differentiation of suprabasal cells and stimulate cell cycle re-entry by interacting with several cellular proteins, including p53 and pRb, respectively [7]. The activity of P105 is determined by regulatory elements, enhancers, and silencers, situated along the viral long control region (LCR) upstream of the promoter [8,9]. Late gene transcription is switched on mostly in keratinocytes of superficial layers undergoing differentiation [10,11].

Orchestrated patterns of viral gene expression during cellular differentiation are intricately controlled by host transcription factors (TFs), underscoring their crucial role in this process [12,13]. To date, numerous cellular TFs that regulate HPV promoter activity by binding at consensus sequences within the LCR have been recognized, including Oct-1, TEF2, NF1, AP-1, YY1, SP1, FOXA1, and GATA-3 [14–18]. However, only a limited number of differentiation-specific TFs implicated in HPV-18 early transcriptional regulation have been identified. For instance, during epithelial differentiation, the levels of YY1, a potent repressor of early gene transcription, and the recruitment of this protein to the HPV-18 LCR are reduced, which results in the disruption of the CTCF-YY1-dependent looping associated with increased LCR chromatin accessibility [19]. Moreover, STAT3 has a critical role in HPV-18 gene expression and episome maintenance in undifferentiated keratinocytes; upon differentiation, STAT3 induces virus genome amplification and late gene expression [20]. Nevertheless, despite the efforts demonstrating the importance of cellular TFs in coupling HPV-18 early gene transcription with differentiation of the infected epithelia, a comprehensive analysis exploring the repertoire of TFs responsible for this link is to our knowledge still lacking

Given the foregoing considerations, we aimed to identify key TFs involved with the finely tuned regulation of the HPV-18 differentiation-dependent life cycle. We initially compared the DNA-binding activity of 345 TFs in nuclear protein extracts obtained from undifferentiated and calcium-induced differentiated HaCaT cells, representing the basal and apical epithelial layers, respectively. We observed differences for 40 TFs, among which we identified, *in silico*, putative binding sites within the HPV-18 LCR for 16. We next selected four TFs to study in more detail: PAX6, HMGB1, NFE2, and FOXI, which have not been previously explored in HPV-18 differentiation-dependent context ChIP assays revealed direct binding of PAX6, HMGB1, and NFE2, but not FOXI1, to the HPV-18 LCR. Taken together our data point towards the involvement of all four TFs evaluated in orchestrating the HPV-18 differentiation-dependent life cycle. Finally, we show that HPV-18 E6 and E7 oncoproteins impair the epithelial differentiation-dependent expression of PAX6, HMGB1, NFE2, and FOXI throughout the stratified squamous epithelium. Advancing the identification of TFs that synchronize viral gene expression with epithelial differentiation provides new insights into the mechanisms involved in the virus life cycle. This progress may also reveal potential new targets for therapeutic intervention.

## Results

### The DNA-binding activity of several transcription factors is shifted upon keratinocyte differentiation

HaCaT are spontaneously immortalized human keratinocytes widely employed to study epithelial differentiation since, upon stimulation, these cells differentiate preserving properties such as differentiation-associated morphogenesis and expression of surface markers [21]. High calcium (>2.8mM) medium triggers HaCaT cell differentiation, which is reversible upon reduction of calcium concentration (<0.3mM) [22–24]. Given our interest in identifying TFs involved in keratinocyte differentiation, we initially validated the HaCaT calcium-induced differentiation model described by Micallef *et al.* (2009) [25]. We evaluated cell morphology and the expression of differentiation makers representative of undifferentiated and differentiated keratinocyte properties. As expected, we observed that cultivating HaCaT cells in high calcium concentration media resulted in morphological alterations with an apparent increase in cell-cell adhesion (**S1 Fig**). Moreover, higher protein levels of the differentiation markers cytokeratin-1 and involucrin were observed in HaCaT cells after five days of culture in high-calcium-containing medium, assuring that this is a suitable and reproducible model to study keratinocyte differentiation-dependent processes.

Next, we compared the DNA-binding activity of a broad range of TFs in nuclear extracts obtained from undifferentiated and Ca^++^-differentiated HaCaT cells representing the basal and suprabasal layers of the stratified squamous epithelium, respectively. The fractionation efficiency of protein extracts was confirmed by immunoblotting once high TATA-binding protein levels were detected in the nuclear fraction, whereas tubulin was undetectable (**S2 Fig**). Nuclear extracts were then incubated with a mix of biotin-labeled DNA oligonucleotide probes containing the consensus binding sequences of 345 different TFs. Protein/DNA complexes formed were separated and hybridized to membranes spotted with the complementary sequence of each TF probe (**S2 Fig and S1 Table**). Relative spot intensities were determined by densitometry (**S2 Table**). **Fig 1** shows the fold change of 40 TFs that demonstrated altered DNA-binding activity upon differentiation, six of which increased upon differentiation (including HIF-1, FOXI, and SP-1). The other thirty-four were downregulated upon exposure to high calcium medium (PAX6, AP-1, PAX8, NFE2, among others).

**Fig 1.**
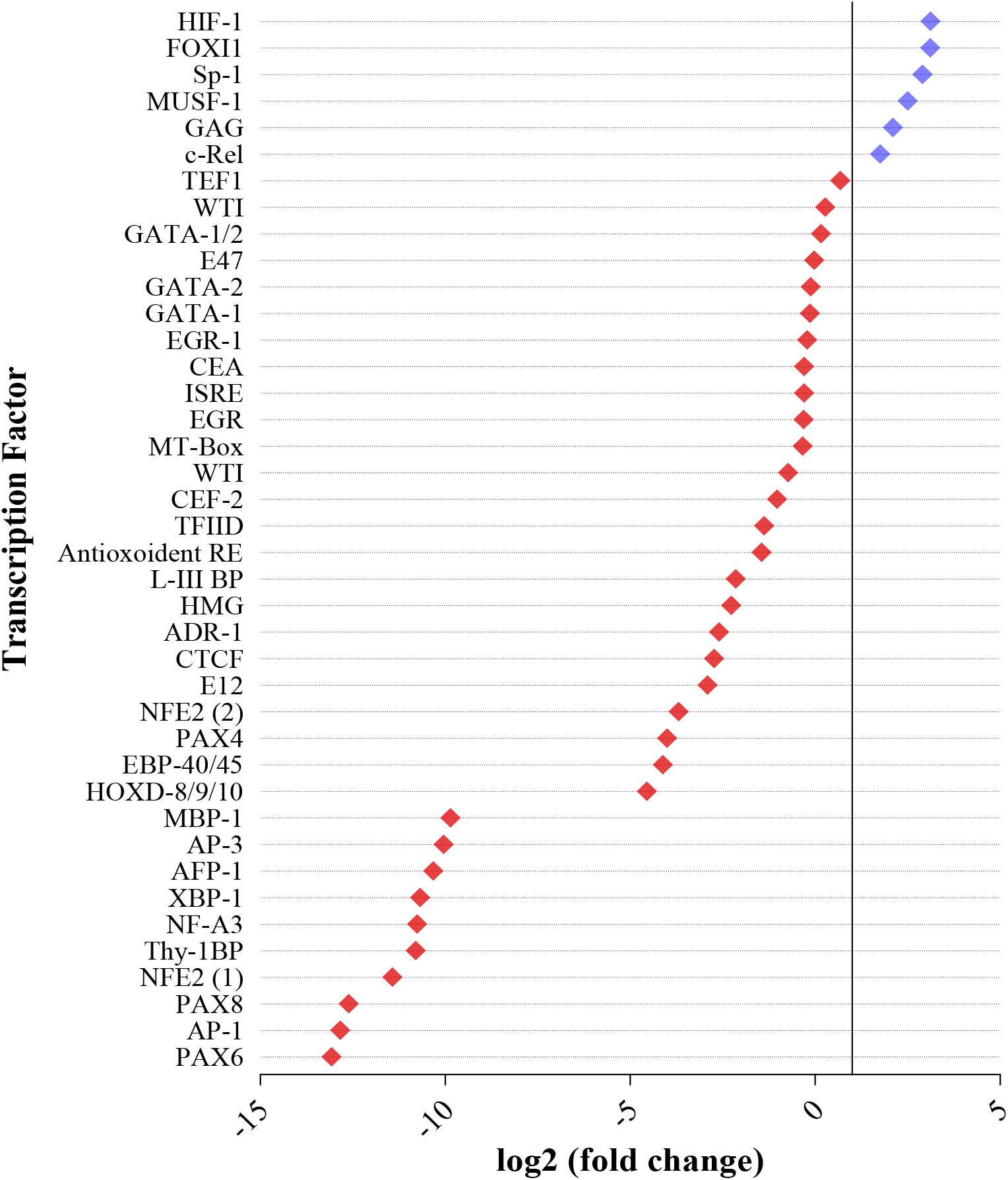
Shift in transcription factors DNA-binding activity induced by HaCaT cell differentiation. Dot intensities of the TransSignal protein/DNA combo array (Panomics, California, USA) membranes from undifferentiated and high-calcium differentiated HaCaT nuclear extracts were compared. The results are shown as log2 fold-change in differentiated versus undifferentiated keratinocytes.

### A subset of differentiation-modulated TFs putatively binds to the HPV-18 LCR

We asked which TFs that exhibit DNA-binding activity during differentiation could directly bind to cis-elements within the HPV-18 LCR. Thus, using the TRANSFAC database, we searched for putative DNA-binding elements within the complete HPV-18 LCR (Genebank X05015) sequence for the 40 TFs which DNA binding activities were altered upon keratinocyte differentiation. **Table 1** shows the genomic position, strand, and nucleotide sequence of the putative binding sites for each TF with the two highest core scores. Core scores were calculated based on the identity between TF motifs and input sequences, indicating the strength of identified matches; values ranged from 0 to 1, with 1 representing an exact match [26]. This analysis successfully identified previously described active AP-1 motifs at nucleotide positions 7613nt and 7795nt of the HPV-18 LCR (core score =1.00) [27,28]. In addition, our analysis revealed putative binding sites for 16 out of the 40 TFs evaluated, including yet to be experimentally validated interactions of AP-3, c-Rel, CTCF, E12, E47, EGR-1, GATA-1, GATA-2, HFH3 (FOXI1), HMG, MT-box, NFE2, PAX4, PAX6, and PAX8.

**Table 1.**
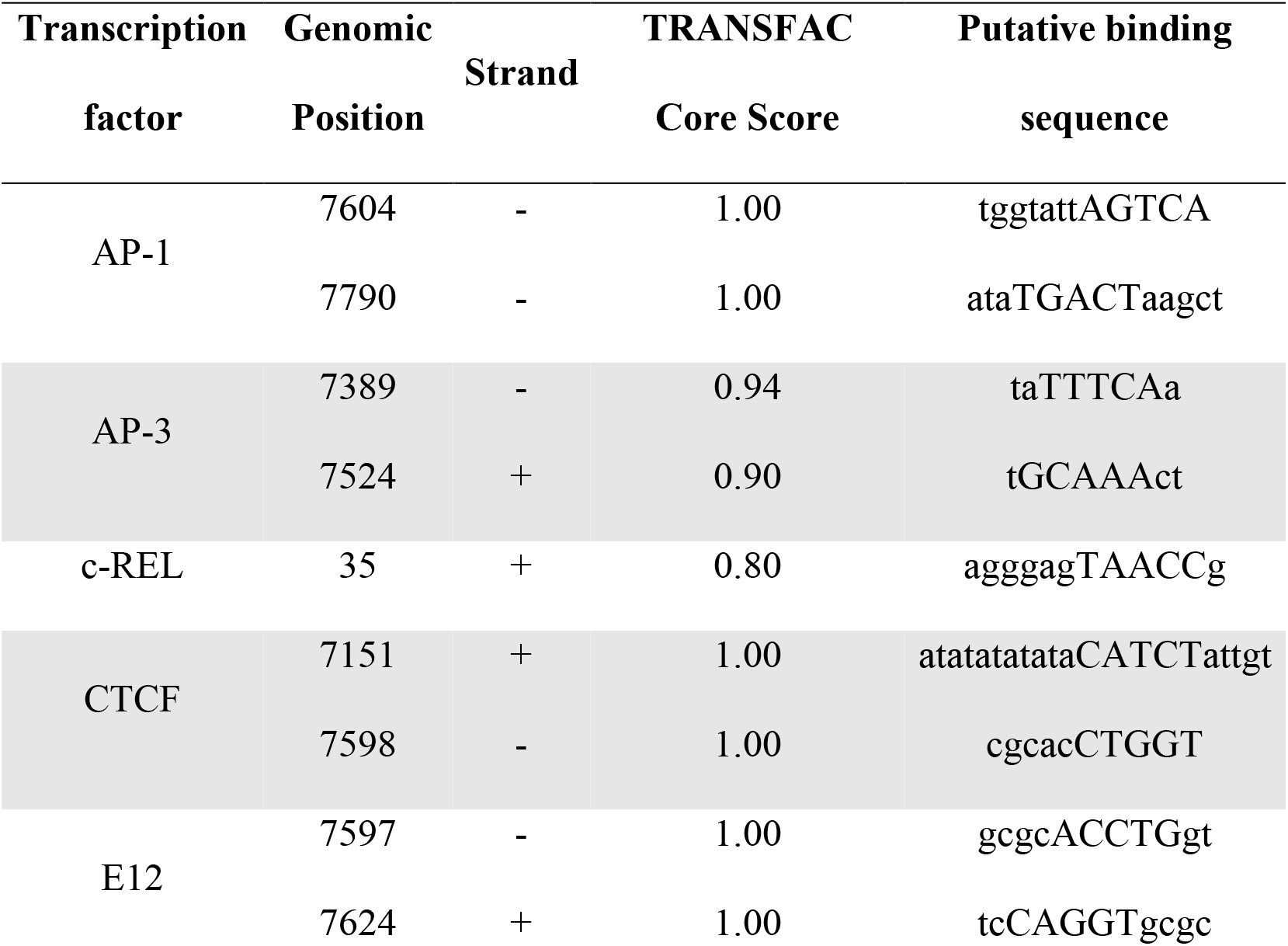
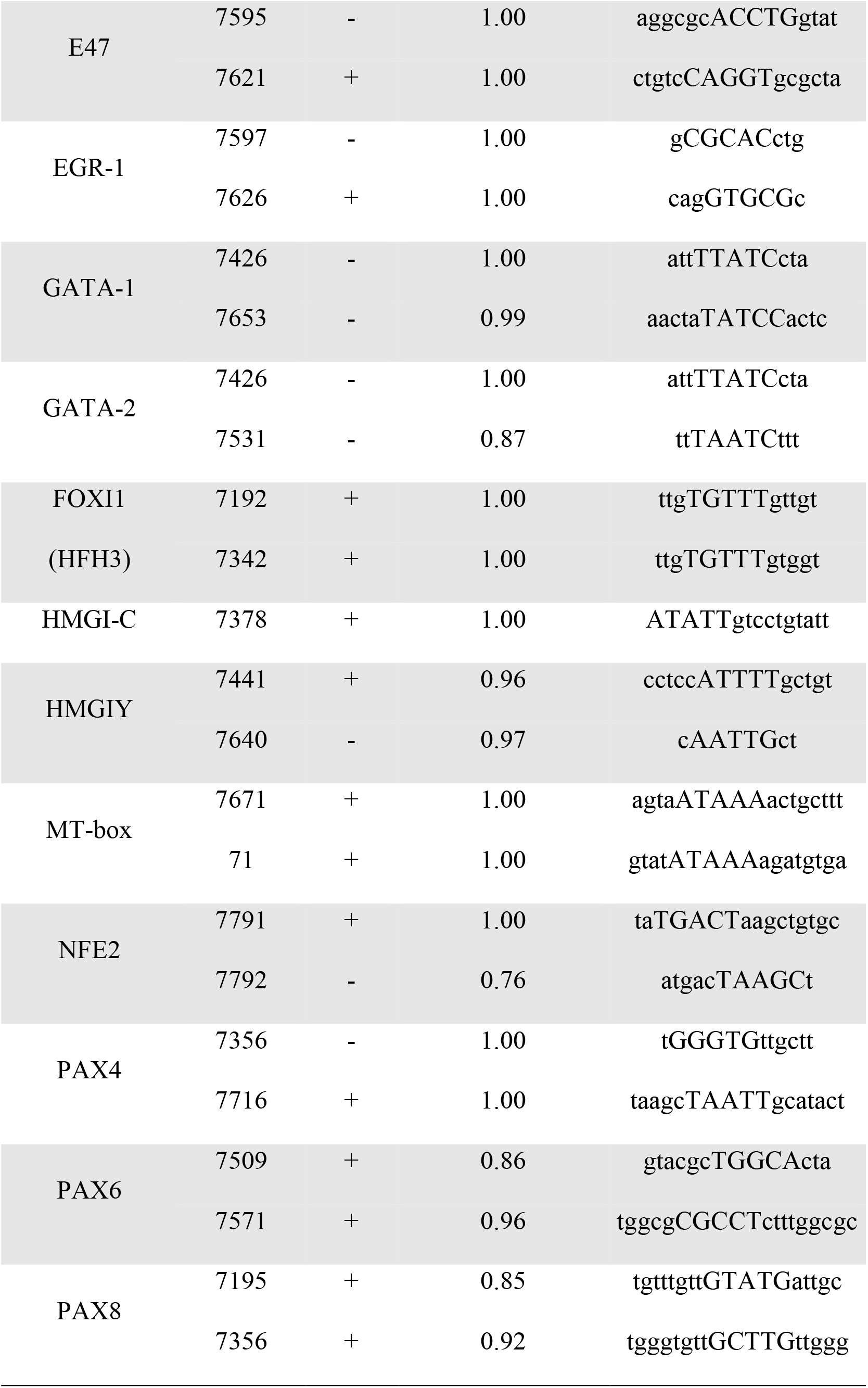
Putative binding sites within the HPV-18 LCR identified *in silico* for selected transcription factors.

We chose to focus our analyses on four TFs not yet described to influence HPVs transcriptional regulation dependent on epidermal differentiation: FOXI (forkhead box I1), for which binding activity increased upon HaCaT differentiation, and PAX6 (paired box protein 6), NFE2 (nuclear factor erythroid 2) and HMGB1 (high-mobility group box 1), for which binding activities decreased upon differentiation.

### PAX6, HMGB1, and NFE2 directly bind to the HPV-18 LCR *in vitro*

To validate the *in silico*-predicted binding of the four selected TFs to the HPV-18 LCR, we used ChIP-qPCR assays. Crosslinked chromatin harvested from 18NCO cells (keratinocytes immortalized with the complete HPV-18 genome) was independently immunoprecipitated with anti-PAX6, -HMGB1, -NFE2, or -FOXI1 antibodies, and enriched fragments were evaluated by qPCR using primer pairs that amplify four separate segments (I-IV) covering the complete LCR sequence. We observed significant binding of PAX6 and HMGB1 to segments III and IV and of NFE2 to segment I of the HPV-18 LCR (**Fig 2A**). None of the fragments analyzed was significantly enriched when FOXI1 was immunoprecipitated. To better assign binding elements of these TFs within the ChIP enriched segments, we employed an integrative approach combining ChIP data with extended *in silico* analysis. Briefly, we searched for PAX6 and HMGB1 binding sites within segments III and IV of the LCR and for NFE2 binding sites within segment I using three distinct data sources: JASPAR [29], HOCOMOCO [30], and TRANSFAC [26] **(S3 Table)**. **Fig 2B** shows the suggested specific binding positions of PAX6, HMGB1, and NFE2 within the HPV-18 LCR, as inferred from enriched segments of ChIP assays and the motifs consistently identified in at least two databases.

**Fig 2.**
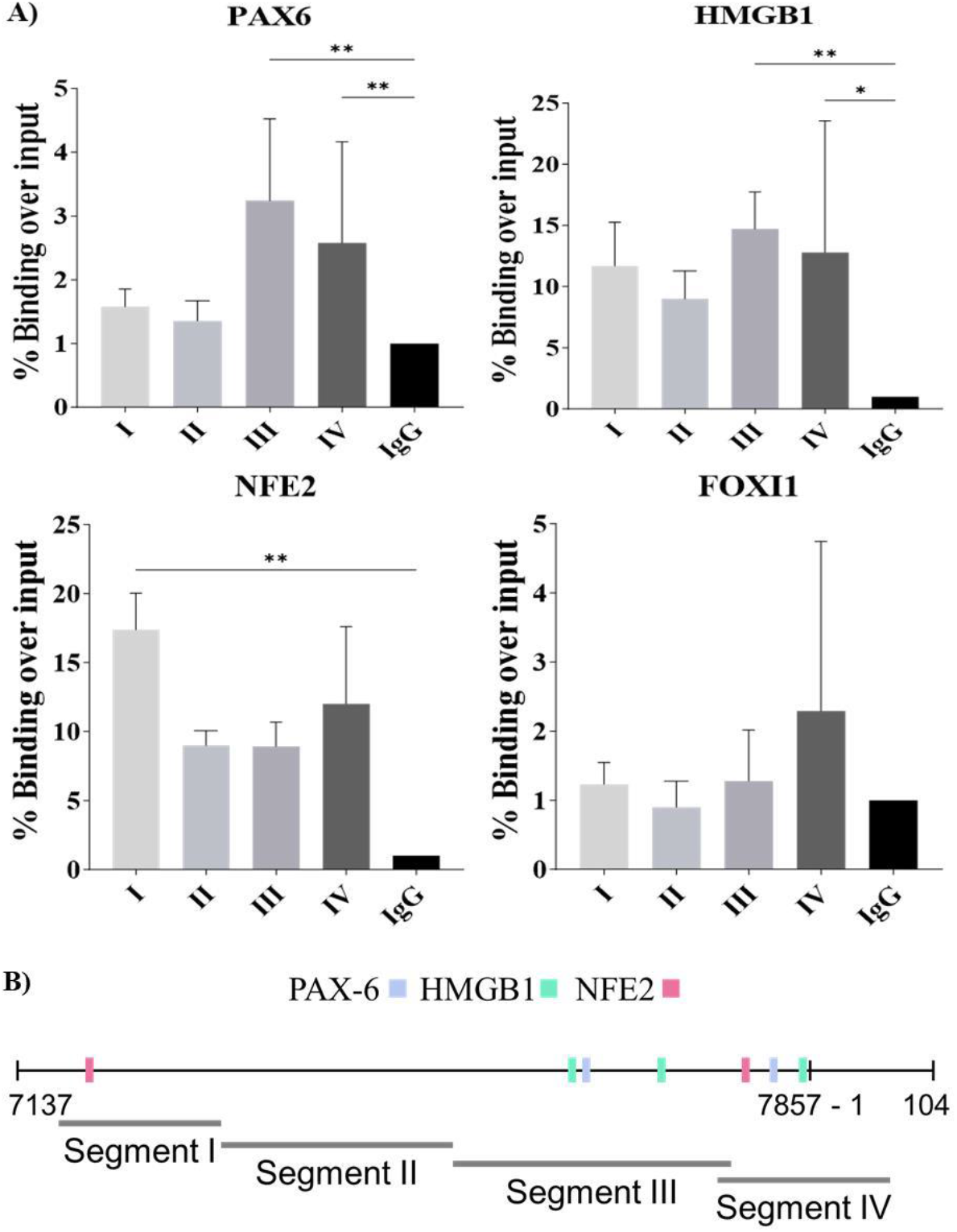
PAX-6, HMGB1, and NF2 bind to cis-elements within the HPV-18 LCR. (A) Direct binding of PAX6, HMGB1, NFE2, and FOXI to the HPV-18 LCR was evaluated by ChIP followed by qPCR using primers that amplify four different segments (I-IV) of the LCR. Data represent binding percentage over input normalized by each assay’s negative control (IgG). The values are represented by the mean ± SD of three independent experiments. (B) Schematic representation of PAX6, HMGB1, and NFE2 presumed binding sites within the HPV-18 LCR based on combined ChIP-qPCR and *in silico* analysis performed using TRANSFAC, HOCOMOCO, and JASPAR databases.

### PAX6, HMGB1, NFE2, and FOXI1 levels decrease upon differentiation

We then assessed whether the differences observed in the DNA binding activity of FOXI1, HMG, NFE2, and PAX6 (**Fig 1**) in differentiated and undifferentiated HaCaT cells was associated with differences in the protein levels of these TFs throughout the stratified epithelium. For this purpose, we performed immunohistochemistry assays in organotypic epithelial raft cultures derived from a pool of passages 0-1 HFK. **Fig 3A** shows intense PAX6 nuclear and cytoplasmic staining in basal layer cells, gradually decreasing towards more differentiated superficial cells. HMGB1 and FOXI1 nuclear staining were observed in all basal cells and sparse cells of suprabasal layers (**Figs 3B** and **3D**). NFE2 expression was confined to the nucleus of cells of the basal layer (**Fig 3C**).

**Fig 3.**
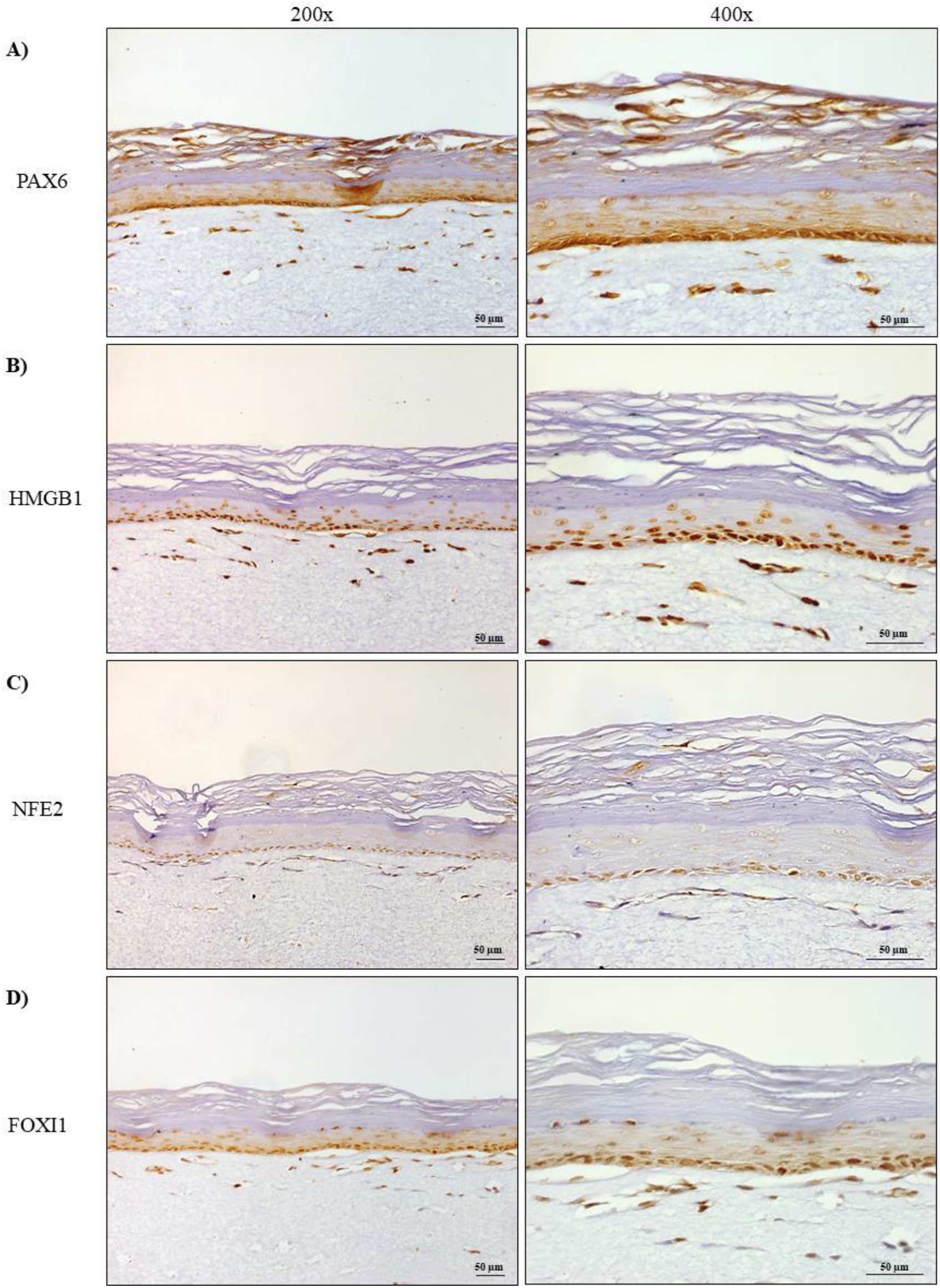
Transcription factors levels and distribution throughout the stratified epithelium. Immunohistochemistry was performed for (A) PAX6, (B) HMGB1, (C) NFE2, and (D) FOXI1 in organotypic cultures derived from a pool of passage 0-1 HFK. Images are representative of two independent experiments. Magnification: 200x (left column) and 400x (right column). Scale bar: 50µm.

### PAX6, HMGB1, NFE2 and FOXI1 impact upon HPV-18 P105 transcriptional activity

Considering that we found the DNA binding activity (**Fig 1**) and the protein levels of PAX6, HMGB1, NFE2, and FOXI1 varied dependent on the differentiation status of the cells (**Fig 3**), and that PAX6, HMGB1 and NFE2 directly bound to the HPV-18 LCR (**Fig 2**), we endeavored to test whether these TFs play a role in synchronization of the viral life cycle by affecting viral early promoter activity. To investigate the impact of PAX6, HMGB1, FOXI1, and NFE2L2 and NFE2L3 (both members of the NFE2 family), upon the transcriptional activity of the HPV-18 P105 promoter, we co-transfected increasing amounts of TF expression plasmids with the LCR-18 Luc reporter plasmid in HPV negative C-33 A cervical cancer cells (**S3 Fig**). We initially performed reporter assays in C-33 A cells because they are easily transfected and cultured, and have been widely used to study HPV transcription [31–33]. We observed that in a dose-response manner, HMGB1 inhibited P105 activity, resulting in transcriptional repression of 63% when the input plasmid was 300ng (**Fig 4B**). Dose dependence was not observed with increased NFE2L2 expression, although HPV-18 early transcription inhibition was observed even in the lowest dose of plasmid transfected (**Fig 4C**). Unexpectedly, PAX-6 and NFE2L3 activated the P105 promoter at lower doses (25ng), while transcriptional repression was observed when higher plasmid quantities were co-transfected (**Figs 4A and 4D**). FOXI was the only TF tested that consistently activated the viral early promoter (**Fig 4E**).

**Fig 4.**
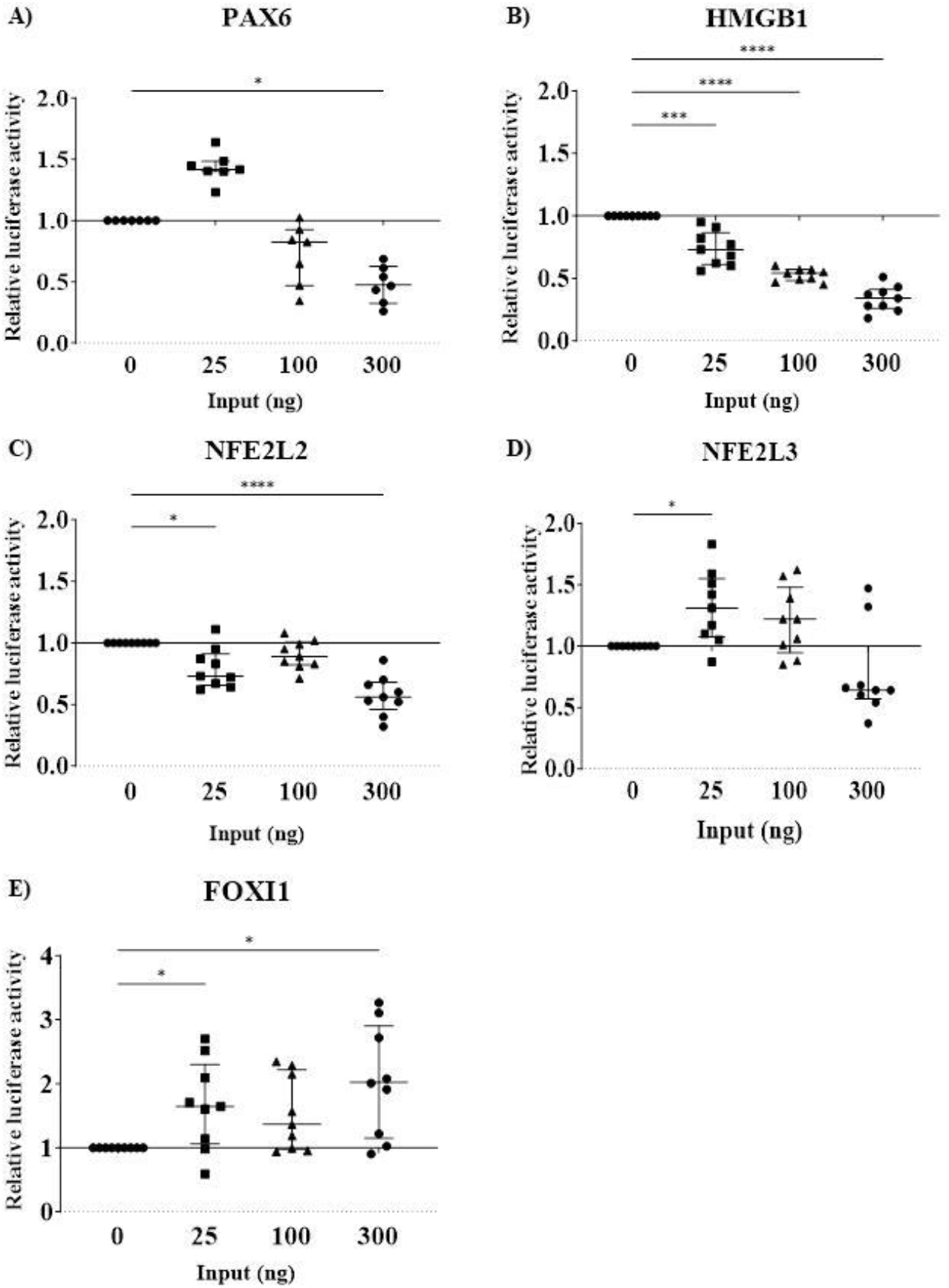
Impact of PAX6, HMGB1, FOXI1, NFE2L2, and NFE2L3 upon the P105 HPV-18 early promoter activity in cervical cancer cells. C-33 A cells were co-transfected with increasing amounts of p-CMV TFs expression plasmids (25, 100, or 300 ng), 100ng of the recombinant reporter vector pGL3-LCR-HPV-18-Luc and 100ng of the pCMV-β-Gal vector. C-33 A cells were co-transfected with the pCMV empty vector (pCMVϕ) as control. After transfection, relative luciferase activities were measured and calculated as the ratio of firefly luciferase/β-galactosidase enzyme activities in the cells and normalized to those of the pCMVϕ control, defined as the reference (value 1). Values below 1 indicate transcription repression, while values above 1 indicate activation of the promoter activity. Data are represented by a median and interquartile range, and each point represents the average of one independent assay conducted in triplicate. Statistical significance: *p<0.05, ***p<0.0005, ****p<0.0001.

### PAX6, HMGB1, NFE2, and FOXI1 regulate HPV-18 early promoter transcriptional activity dependent on the keratinocyte differentiation state

We asked if the impact of these TFs upon the HPV-18 P105 promoter activity is associated with the differentiation status of HaCaT cells. Initially, we determined the basal early promoter activity in undifferentiated compared to differentiated HaCaT cells, and as expected significant downregulation of P105 activity was observed upon cellular differentiation (**Fig 5A**). We next evaluated the effect of TFs overexpression under these conditions. We observed that PAX6 and HMGB1 repressed P105 in both undifferentiated and differentiated HaCaT cells. However, whereas transcription repression by PAX6 was significantly higher upon HaCaT differentiation (0.74 versus 0.57, p=0.023), HMGB1 repression was relieved when cells underwent differentiation (0.37 versus 0.53, p=0.028) (**Figs 5B** and **5C**). Notably, both NFE2 family members, NFE2L2 and NFE2L3, activated HPV-18 early transcription in undifferentiated keratinocytes, while upon induction of differentiation, repression of P105 activity was observed (**Figs 5D** and **5E**). Finally, we corroborated the role of FOXI1 as an HPV-18 transcriptional activator in keratinocytes and observed that induction of transcription was even higher in undifferentiated keratinocytes compared to differentiated keratinocytes (4.57 versus 2.47, p= 0.0003) (**Fig 5F**).

**Fig 5.**
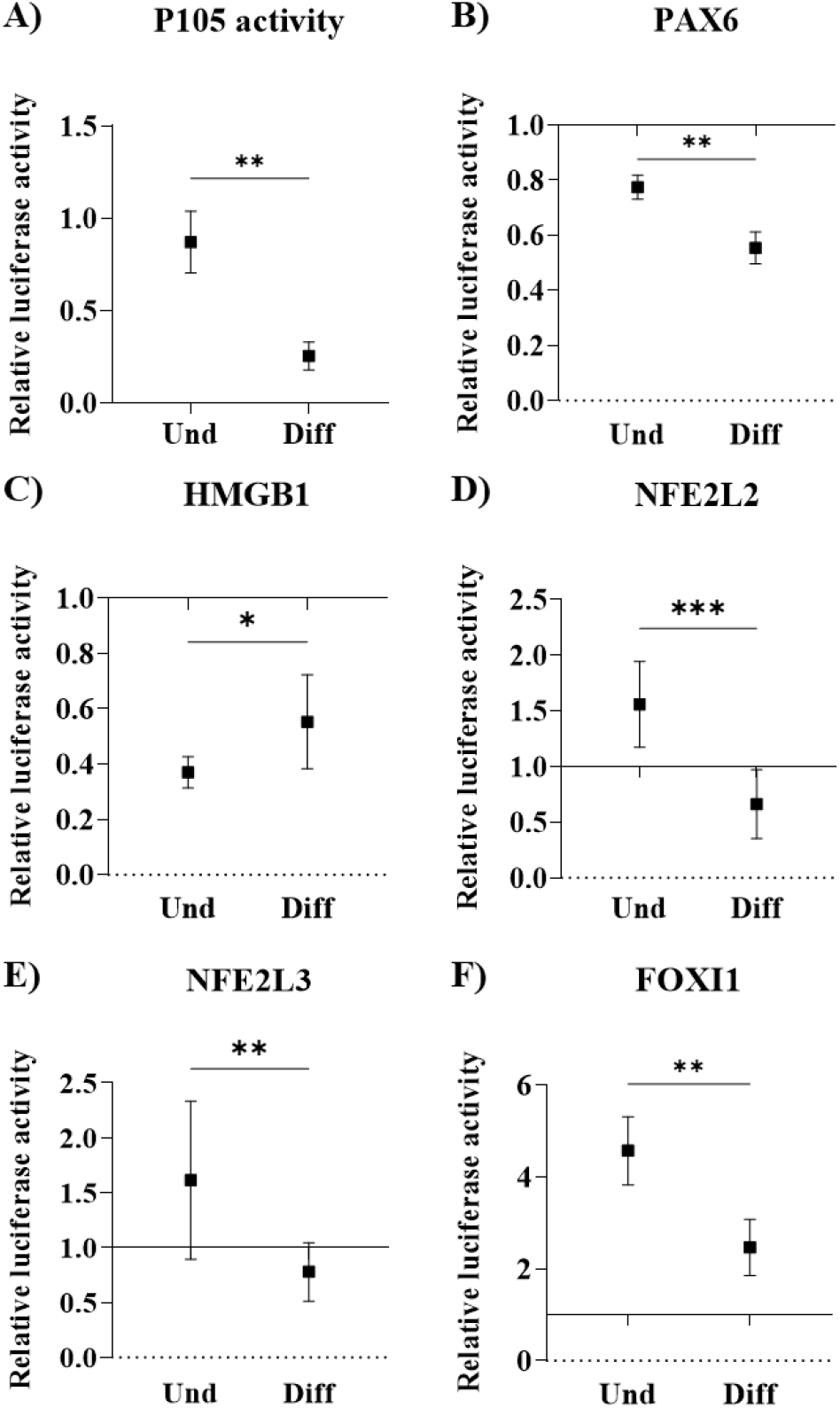
Impact of PAX6, HMGB1, FOXI1, NFE2L2, and NFE2L3 upon HPV-18 transcriptional activity in undifferentiated and differentiated HaCaT cells. HaCaT cells were maintained in media without calcium – undifferentiated (Und) – or in high calcium concentration media for 5 days – differentiated (Diff). At day 3, HaCaT cells in both conditions were co-transfected with either a p-CMV-empty vector or p-CMV TFs expression plasmid plus the pGL3-LCR HPV-18-Luc recombinant reporter vector and the pCMV-β-Gal vector. After 48 hours, completing 5 days, total protein was extracted and the reporter assays were performed. Values are represented by the mean and standard deviation of at least five independent experiments conducted in triplicate. Statistical significance: ns, not significant, *p<0.05, **p<0.005, ***p<0.0005.

### PAX6, HMGB1, NFE2, and FOXI1 expression throughout the stratified epithelium is impaired by HPV-18 E6 and E7 oncoproteins

High-risk HPVs inhibit cell differentiation and induce carcinogenesis by interacting with a plethora of host cellular proteins, among which the most studied associations are of E6 and E7 with p53 and pRb, respectively [34,35]. Abnormal expression of cellular TFs could contribute to HPV persistence by modulating viral transcription and consequently affecting E6 and E7 levels. Thus, we compared PAX6, HMGB1, NFE2, and FOXI1 levels of parental HFK and HFK18E6E7 cells, which are HFK immortalized with HPV18 E6 and E7. RT-qPCR quantification of the TF mRNA levels in undifferentiated monolayer cells revealed that, except for NFE2L2, all transcripts were significantly higher in HPV-18 E6 and E7 immortalized HFK (**Fig 6A**).

**Fig 6.**
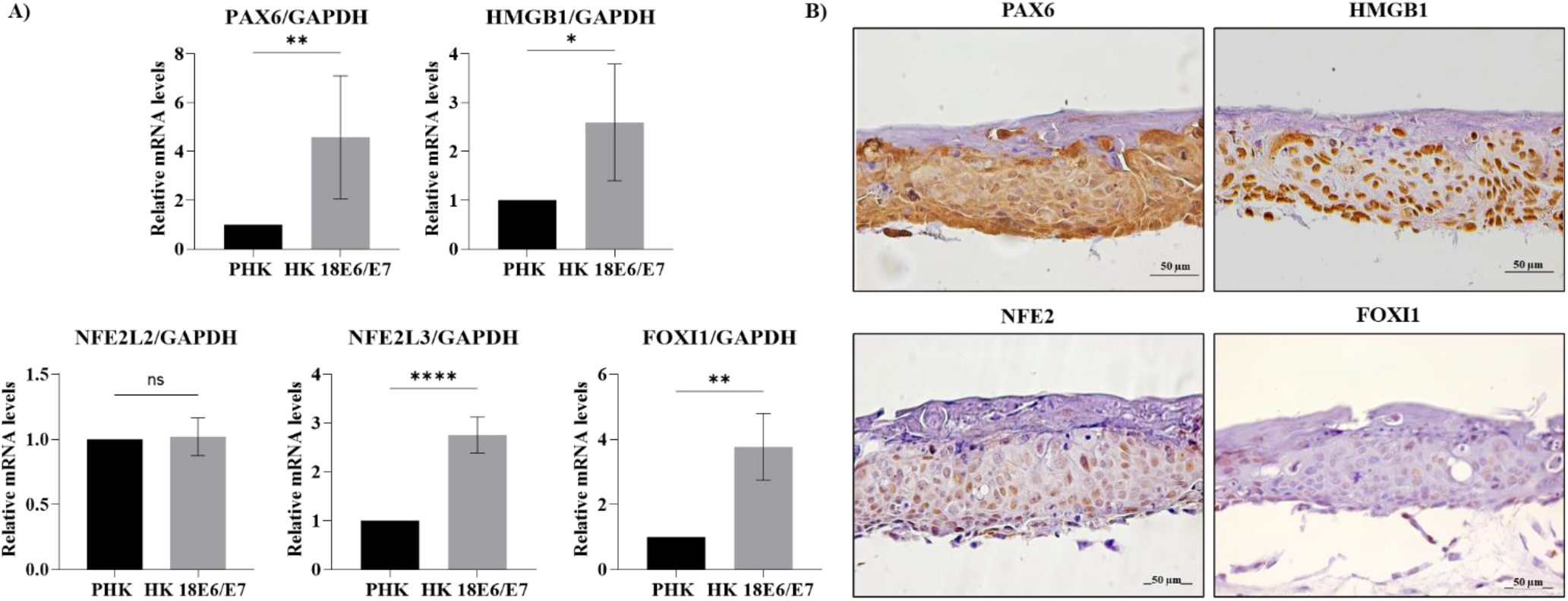
HPV-18 E6 and E7 impairs PAX6, HMGB1, NFE2, and FOXI1 expression throughout the stratified epithelium. (A) Total RNA was extracted from monolayer cultures of primary keratinocytes (HFK) and keratinocytes immortalized with HPV-18 E6 and E7 (HFK18E6/E7). TFs mRNA levels were measured by qPCR and normalized by GAPDH levels. Data are represented by the mean and standard deviation of at least three independent experiments. Statistical significance: ns, not significant, *p<0.05, **p<0.005, ****p<0.0001. (B) Immunohistochemical analyses of organotypic cultures derived from HFK18E6E7 were performed for PAX6, HMGB1, NFE2, and FOXI1. Images are representative of two independent experiments. Original magnification: 200x. Scale bar: 50µm.

Finally, we compared PAX6, HMGB1, NFE2, and FOXI1 protein levels and distribution in organotypic epithelial raft cultures obtained from HFK (**Fig 4**) and HFK18E6E7 cells (**Fig 6B**). In contrast to the organized epithelial stratification resembling normal skin observed in organotypic raft cultures derived from HFK, tissues obtained from HPV-18 E6/E7 immortalized HFK exhibited hyperplasia, and an aberrant stratification that hindered the discrimination between basal and suprabasal layers. Overall, we observed that the TFs analyzed, whereas protein expression was more limited to the basal layers in non-immortalized parental HFK-derived rafts, in HFK18E6/E7-derived rafts protein staining was more intense and widespread across all cell layers, except for FOXI1 which presented lower levels in HFK18E6/E7-derived rafts basal layers compared to HFK-derived rafts, although we observe a more diffuse expression across all layers (**Fig 6B**).

## Discussion

HPV relies on keratinocyte differentiation to complete its life cycle highlighting the importance of exploring the molecular mechanisms that govern this synchronization [36,37]. The interplay between binding elements for host cell TFs within the LCR and the finely regulated temporal and spatial expression patterns of these TFs throughout layers of the stratified epithelium are pivotal players in this process [12,38]. It has been, for example, reported that AP-1, Sp1, Sp3, C/EBPβ, and STAT are expressed in a coordinated manner during epithelial differentiation [20,39–41]. In this study, we compared the transcriptional profile of undifferentiated and calcium-induced differentiated keratinocytes to further identify TFs implicated in HPV differentiation-dependent transcriptional regulation.

We initially used a comprehensive approach to evaluate simultaneously the DNA binding activities of 345 TFs. This sophisticated methodology has been previously used to identify changes in the levels of cellular TFs after methylcellulose-induced differentiation of 20863 cells (a subclone of the W12 cell line derived from an HPV-16 positive low-grade squamous intraepithelial lesion) [38]. Herein, we opted to use HaCaT cells which more closely resemble the cellular environment of non-infected epithelial cells, and further took advantage of a well-established keratinocyte differentiation model [25]. Interestingly, we identified a distinct set of TFs regulated by differentiation in comparison to the previous study conducted in 20863 cells. Among the 345 consensus binding sequences included in the array (**S1 Table**), we observed changes in the binding of 40 TFs after high calcium exposure (**Fig 1, S2 Fig**), among which HIF-1 (3.12-log2FoldChange) and FOXI1 (3.11-log2FoldChange) were the most upregulated, while PAX6 (13.07-log2FoldChange) and AP-1 (12.84-log2FoldChange) were the most downregulated (**S2 Table**).

To investigate TFs that could affect HPV-18 transcriptional regulation, we first evaluated *in silico* which of the 40 TFs identified could potentially bind to the HPV-18 LCR using the TRANSFAC database. Putative binding sites for 16 TFs were revealed (**Table 1**), most of which are located within the central segment of the LCR where an epithelial-specific enhancer is located [42,43]. Several of these TFs have not been previously associated with the differentiation-dependent HPV-18 transcriptional regulation including AP-3, FOXI1 (HFH3), HMG, NFE2, and members of the PAX and GATA families’ members. In this study, driven by insights from the existing literature and the commercial availability of ChIP-grade antibodies, we focused analysis on four TFs: PAX6, HMGB1, NFE2, and FOXI.

ChIP-qPCR assays, conducted in HPV-18 full-genome harboring 18NCO cells, confirmed direct binding of PAX6, HMGB1, and NFE2 to the LCR (**Fig 2A and S4 Table**). However, once these assays validated TFs binding to LCR segments of approximately 200bp, we extended the *in silico* analysis using two additional databases (HOCOMOCO and JASPAR) in order to refine TFs putative binding positions within the enriched fragments (**Fig 2B**). Even though the combined *in silico* and *in vitro* analyses strongly support the direct binding of PAX6, HMGB1, and NFE2 to the HPV-18 LCR, we acknowledge the limitation of our study in precisely determining the binding sites position. Such determination would require extensive and laborious mutagenesis analysis, which, however, deviates from the main goal of our study. Lastly, ChIP assays did not detect significant binding of FOXI to the HPV-18 LCR.

To evaluate the association of PAX6, HMGB1, NFE2, and FOXI1 expression with the epithelial differentiation program, we analyzed the levels of these proteins in raft cultures grown from HFKs [44]. Overall, we observed that all four TFs levels are higher in undifferentiated basal layer cells compared to differentiated cells of the upper layers (**Fig 3 and S4 Table**). Thus, the decreased DNA binding activities of PAX6, HMGB1, and NFE2 after exposure to high calcium (**Fig 1**) may be associated with higher levels of these proteins in undifferentiated cells. Unexpectedly, though FOXI1 DNA binding activity was upregulated after differentiation (**Fig 1**), FOXI1 IHQ staining was mostly confined to undifferentiated basal cells (**Fig 3**), with only a minority of differentiated cells expressing this TF. Despite the diverse biological functions of PAX6, HMGB1, and NFE2, data concerning the involvement of these TFs in the context of keratinocyte differentiation is still very limited. To our knowledge, the link between HMGB1 and cell differentiation was evidenced in only one study which demonstrated that the functional ablation of secreted HMGB1 results in aberrant HaCaT differentiation [45]. Additionally, NFE2 was shown in several studies to activate antioxidant enzymes to protect skin keratinocytes against oxidative damage during differentiation [46–48]. Finally, the involvement of PAX6 and FOXI1 in differentiation has been evaluated only in cell types other than genital and cutaneous keratinocytes; while it was demonstrated that PAX6 plays a pivotal role in directing the differentiation and cellular fate of corneal epithelial cells [46,49–51], FOXI1 was proven important for renal epithelial cells differentiation [52,53].

To further explore the role of PAX6, HMGB1, NFE2, and FOXI in linking the HPV life cycle to the epithelial differentiation program, we evaluated the impact of these TFs upon viral early promoter activity. We initially explored, in a dose-response manner, the influence of these TFs upon HPV-18 P105 activity in C-33 A cells and observed that in the higher plasmid input tested PAX6, HMGB1, NFE2 significantly inhibited early transcription, in contrast to the positive stimulus induced by FOXI (**Fig 4**). Next, to evaluate the interdependence of these TFs’ effect with the cell differentiation status, we conducted reporter assays in both undifferentiated and differentiated HaCaT cells. Unfortunately, carrying out protein-DNA binding and luciferase reporter assays in differentiated HFKs is unfeasible as these cells are not viable post-differentiation [54,55].

Taken together, our data points towards the involvement of all four TFs evaluated in orchestrating the HPV-18 differentiation-dependent life cycle. FOXIl activated HPV-18 transcription in both C-33 A and HaCaT cells, exhibiting a significantly more pronounced impact in undifferentiated cells (**Fig 5F**) where this TF is mostly expressed (**Fig 3D**), and potentially contributing to the higher P105 early promoter activity observed in these cells (**Fig 5A**). Although higher FOXI1 DNA-binding activity was observed in differentiated cells (**Fig 1**), we believe this TF indirectly activates HPV-18 transcription as no FOXI1-LCR binding was uncovered by ChIP assays (**Fig 2A**). Accordingly, previous studies have highlighted the importance of partner proteins and co-factors recruitment for the activity of forkhead TF family members [56,57]. NFE2 protein members also seem to contribute to HPV-18 early promoter activation in undifferentiated cells (**Fig 3E**), however by directly binding to the LCR (**Fig 2A**). On the other hand, PAX6 and HMGB1 significantly inhibited P105 in undifferentiated HaCaT cells (**Fig 5**) in consonance with higher DNA-binding activity and protein levels in these cells (**Fig 1 and Fig 3A and B**). Although this observation seems contradictory to the fact that basal early promoter activity is higher in HaCaT undifferentiated cells in comparison to the differentiated counterpart (**Fig 5A**), it is noteworthy that transcription is a dynamic process resulting from the stoichiometric balance of a plethora of enhancer and inhibitory TFs expression and activity [58,59]. Importantly, this equilibrium may be relevant for keeping early protein expression to a minimum to favor HPV escaping from immune recognition [60,61]. Lastly, though HMGB1 and PAX6-mediated P105 inhibition was significantly alleviated and enhanced, respectively (**Fig 5B and C**), upon cell differentiation, we should not further elaborate on this finding as in normal epithelial both protein levels are very low or inexistant in suprabasal layer cells (**Fig 3A and B**).

It is well described that high-risk HPV E6 and E7 oncoproteins modulate the activity of various cellular proteins involved in cell proliferation and differentiation, including TFs [7,62,63]. For instance, members of the AP-1 complex, known to play a seminal role in inducing HPV transcription in undifferentiated basal cells [64,65], were shown to be expressed predominantly in proliferating basal cells in normal cervix and low-grade cervical intraepithelial neoplasia (CIN) samples, in contrast to high-grade CIN and invasive cervical cancer in which expression was detected across all epithelial layers [66]. Thus, we speculated whether HPV-18 E6 and E7 oncoproteins influence PAX6, HMGB1, NFE2, and FOXI expression and localization. Interestingly, in the background of raft cultures obtained from HPV-18 E6 and E7 immortalized cells [44], we observed, in general, the high levels of all TFs evaluated throughout the entire thickness of the epithelia (**Fig 6**), in contrast to the normal epithelial where expression of these proteins is mostly limited to basal layers cells (**Fig 3**). While the data presented here is preliminary and further studies are warranted, it suggests a potential relevance of PAX6, HMGB1, NFE2, and FOXIl in this scenario where epithelial differentiation is disturbed and proliferation must be maintained, such as observed in HPV immortalized cells. Moreover, these FTs may also directly contribute to the dysregulation of several pathways implicated in HPV-induced carcinogenesis. In fact, previous studies have shown that abnormal expression of HMGB1 enhances the proliferative and tumorigenic abilities of cervical cancer cell lines and is linked to poor prognosis and metastasis among women diagnosed with cervical cancer. [67–69]. Additionally, targeting the NFE2L2 pathway in cervical and endometrial cancers was shown to improve response to chemotherapy and to suppress cell proliferation [70–72]. Although PAX6, NFE2L3, and FOXI have not yet been explored in the context of HPV-related malignancies, the role of these TFs in other cancers has been reported. For instance, PAX6 was shown to support cell proliferation and tumor growth in lung and breast cancer [73–75]. Additionally, NFE2L3 was associated with several cancer hallmarks, including in promoting the proliferation of cells obtained from multiple cancer types [76–78], and FOXIl was shown to influence breast cancer growth and metastasis and potentially serve as a biomarker for renal cancer [79–81].

In conclusion, we describe the involvement of PAX6, HMGB1, NFE2, and FOXIl in coupling HPV-18 transcriptional regulation with the epithelia differentiation program, a key event for the productive HPV life cycle. Furthermore, HPV-18 E6/E7 impacted upon abnormal expression of these TFs, which may, be also relevant for HPV-driven carcinogenesis. The identification of new cellular targets involved in regulating the differentiation-dependent HPV life cycle and associated cellular transformation may contribute to unveiling novel therapeutic targets for HPV-related malignancies.

## Materials & Methods

### Cell Culture

Undifferentiated HaCaT cells (BCRJ, Rio de Janeiro, Brazil), primary human foreskin keratinocytes (HFK) (Lonza, Basel, Switzerland), and HPV-18 E6/E7 immortalized HFKs (HFK18E6E7) [44] were cultured in KSFM (Keratinocyte Serum Free Medium, Life Technologies, California, USA) supplemented with 0.2 ng/ml of EGF (epidermal growth factor) and 30 µg/ml of BPE (bovine pituitary extract). To induce HaCaT cell differentiation, calcium chloride (Sigma-Aldrich, Missouri, USA) was added to the culture medium at a final concentration of 2.8 mM for up to 5 days. HFK18E6E7 were previously generated with complete HPV-18 E6/E7 genes of A1 lineage variants cloned within the pLNSX retroviral vector (GenBank M28246.1) [44]. The 18NCO cell line (human keratinocytes immortalized with the complete HPV-18 genome) was provided by R. Schlegel, Georgetown University Medical Center, Washington, DC [82], and was grown in “3 + 1” medium, consisting of 3 parts of KSFM supplemented with 0.2 ng/ml of EGF and 30 µg/ml of BPE, and 1 part of DMEM (Dulbecco’s Modified Eagle Medium, Gibco/ThermoFisher, Massachusetts, USA) supplemented with 10% fetal bovine serum (FBS). The HPV-negative cervical carcinoma cell line C-33 A (ATCC HTB-31) was maintained in Dulbecco’s modified Eagle medium (DMEM) supplemented with 10% FBS. All cells were maintained at 37°C and 5% CO_2_.

### Immunoblotting

Cell lysates were obtained using RIPA buffer (20mM Tris-HCl pH=7.5, 150mM NaCl, 0.5% sodium deoxycholate, 0.1% SDS, 1% NP40) containing protease and phosphatase inhibitors (Roche, Basel, Switzerland). 40μg of each protein extract (total or nuclear protein extract) were fractionated by SDS–PAGE and transferred to PVDF membranes (GE Healthcare Life Sciences, Buckinghamshire, UK). These were blocked in 5% nonfat dry milk in TBS-T (20mM Tris-HCl pH=7.5, 150nM NaCl, 0.1% Tween20) for one hour and then immunoblotted. Membranes were probed with the following primary antibodies: anti-cytokeratin 1 (ab193652, Abcam, Massachusetts, EUA, 1:500); anti-involucrin (ab53112, Abcam, Massachusetts, EUA, 1:300); anti-GAPDH (G8795, Sigma-Aldrich, Missouri, USA, 1:2000), anti-TATA binding protein (ab818, Abcam, Massachusetts, EUA, 1:500) or anti-tubulin (ab56676, Abcam, Massachusetts, EUA, 1:5000). Blots were developed using the ECL Detection Reagent (GE Healthcare Life Sciences), images were acquired on a ImageQuant LAS 4000 (GE Healthcare Life Sciences) and analyzed using Image J 1.52p (National Institute of Health, Bethesda, MD, USA).

### Nuclear Protein Extraction and Protein/DNA array hybridization

Nuclear proteins from undifferentiated and differentiated HaCaT cells were obtained using the Panomics nuclear extraction kit (Panomics, California, USA). Next, the DNA binding activity of 345 different TFs was evaluated in these extracts using the TransSignal protein/DNA combo array (MA1215, Panomics, California, USA) according to the manufacturer’s instructions. Membrane signals were developed and quantified using the ImageQuant ™ TL 8.1 software (GE Healthcare Life Sciences, Buckinghamshire, England).

### *In silico* search for TF Binding Sites within the LCR

We searched for putative TF binding sites within the HPV-18 LCR using TRANSFAC 2.0 database (https://genexplain.com/transfac/) [26], HOmo sapiens COmprehensive MOdel COllection (HOCOMOCO) v11 (https://hocomoco11.autosome.org/) [30], and JASPAR 10^th^ release (https://jaspar.elixir.no/analysis) [29].

### Organotypic Epithelial (Raft) Tissue Cultures and Immunohistochemistry

We accessed protein levels and distribution of selected TFs using previously grown epithelial raft cultures obtained from two different pools of HFK immortalized or not by HPV-18 E6/E7 from the A lineage (named here as HFK18E6E7) [44]. Parental HFKs at *p*0 and HFK18E6E7 were submitted to epithelial raft cultures, as described [83]. Briefly, parental HFK and HFK18E6E7 were seeded on top dermal equivalents (2× 10^5^ cells/equivalent) composed of rat tail type 1 collagen (Corning Inc., Corning, NY, USA) and 3T3-J2 fibroblasts. After 24h, the rafts were transferred to the medium–air interface and maintained for 9 days to allow cell growth and tissue stratification. We performed two independent raft experiments with at least six replicates. Epithelia were fixed in formaldehyde 2%, paraffin-embedded, and tissue sections obtained for histological analysis or immunohistochemistry (IHC). After deparaffinization in xylene and rehydration in alcohol, antigen retrieval was performed by incubation in boiling 10 mM citrate buffer, pH=6.0, for 10 minutes. Slides were incubated with anti-PAX6 (ab5790, Abcam, Cambridge, England, 1:400), anti-HMGB1 (ab18256, Abcam, Cambridge, England, 1:2000), anti-NFE2 (ab140598, Abcam, Cambridge, England, 1:400), or anti-FOXI1 (PA530031, Life Technologies, California, USA, 1:250), for 18 hours at 4°C. Reactions were developed using the Polymer Detection System (NovolinkTM Max Polymer Detection Systems, Leica Biosystems, Newcastle, UK) according to the manufacturer’s instructions. Positive reactions were visualized with a cocktail containing peroxidase-conjugated anti-mouse or anti-rabbit secondary antibodies in the presence of DAB (3,3’-diaminobenzidine tetrahydrochloride) resulting in a brown precipitate.

### Chromatin Immunoprecipitation Assays

18NCO cell chromatin fragmentation was performed by sonication with 15 pulses of 30 seconds using the Ultrasonic Bath (Cristófoli, Paraná, Brazil) with 45 seconds of incubation in ice between pulses. The size of the produced fragments was verified by 1% agarose gel electrophoresis. To access in vivo DNA binding of TFs we used the MAGnify Chromatin Immunoprecipitation System (ThermoFisher, Massachusetts, USA) following the manufacturer’s instructions. Briefly, 1µg of chromatin (corresponding to 10^5^ cells) was incubated overnight with 0.8μg of each antibody (anti-PAX6, anti-HMGB1, anti-FOXI, and anti-NFE2, Abcam, Cambridge, England), at 4°C with moderate agitation. The binding percentage over input was calculated as follows:

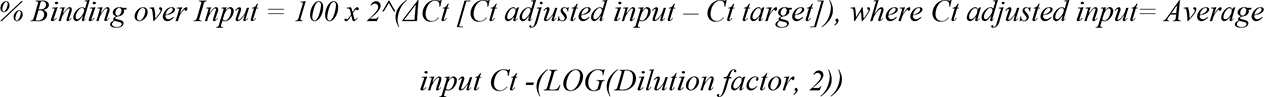

The data are shown by mean and standard deviation of fold change relative to the non-immune IgG sample of three independent experiments.

### Reverse transcription and quantitative PCR

Total RNA was extracted using Trizol (Life Technologies, California, USA) according to the manufacturer’s protocols. TFs mRNA levels were evaluated by reverse transcription followed by real-time quantitative PCR (RT-qPCR) using the GoTaq 1-Step RT-qPCR System (Promega, Madison, USA) and primers listed in **S5 Table**. For ChIP analysis, recovered DNAs were subjected to a real-time polymerase chain reaction (qPCR) using specific primers that amplify four different segments of the HPV-18 LCR (**S6 Table**). The GoTaq qPCR Master Mix (Promega, Madison, USA) was used following the manufacturer’s instructions. Reactions were performed on a 7500 Real-Time PCR System (Applied Biosystems, Foster City, EUA).

### Transient co-transfection reporter assays

C-33 A cells were seeded into 96-well plates and co-transfected with 100ng of the previously constructed recombinant vector pGL3-HPV-18 LCR-Luc [32], 100ng of the pCMV-β-Gal vector, which was used as an internal control of transfection efficiency, and increasing amounts (25, 100 and 300ng) of cDNA-containing expression plasmids from selected TFs (pCMV-PAX6, pCMV-HMGB1, p-CMV-NFE2L2, p-CMV-NFE2L3 or p-CMV-FOXI) (Origene, Rockville, USA), or with the pCMV empty vector (pCMVϕ). Furthermore, undifferentiated or differentiated HaCaT cells were seeded into 6-well plates and co-transfected with 1µg pGL3-Basic HPV-18 LCR-Luc together with 1μg of the pCMV-β-Gal vector, and 500ng of the selected TFs expression vectors. The Fugene 6 reagent (Promega, Madison, USA) was used at a 3:1 rate, following the manufacturer’s instructions. After 48 hours of transfection, luciferase and β-galactosidase activities were measured using the Promega Luciferase Assay System (Promega, Madison, USA) and the Promega β-Galactosidase Enzyme Assay System (Promega, Madison, USA), respectively. Six to nine independent experiments were performed according to standard deviations.

### Statistical analysis

Statistical significance was determined using the student t-test or one-way ANOVA (**Fig 4**) with Prism7 (GraphPad). The levels of statistical significance for each experiment (*, p<0.05; **, p<0.005; ***, p<0.0005; ****, p<0.0001; ns, not significant) are indicated in the corresponding figures. Error bars in the graphs represent the standard deviation.

## Acknowledgments

We are grateful to Dr. Enrique Mario Boccardo, Dr. Vanesca de Souza Lino, and Dr. Emily Montosa Nunes for the donation of HFK and HFK18E6E7-derived rafts. We are grateful to Dr Michelle A Ozbun for kindly collaborating with insightful comments.

## Supporting information captions

**S1 Fig. Differentiation of HaCaT cells induced by high calcium concentration containing media.** Undifferentiated HaCaT cells (day 0) were cultured for 5 days in a media containing 2.8mM of calcium to induce cell differentiation (day 5). Phase contrast images show the morphological differences between (A) undifferentiated and (B) differentiated HaCaT cells. Total protein extracts of HaCaT cells in both conditions were analyzed by Western blot. Representative immunoblots of (C) cytokeratin-1 and (D) involucrin. GAPDH was used as a loading control. (E) and (F) Relative protein levels were obtained from densitometric analysis of immunoblots. Data from two independent experiments were collected and plotted showing the mean and ±SEM. Statistical analysis was carried out using Student’s t-test to obtain significant values. * p < 0.05, ** p < 0.01.

**S2 Fig. Evaluation of DNA-binding activity of 345 TFs in nuclear extracts obtained from undifferentiated and differentiated HaCaT cells.** (A) Nuclear extract of HaCaT cells in both conditions (U= undifferentiated; D= differentiated) were subjected to immunoblotting to confirm the fractionation effectiveness. The level of 345 TFs in (B) undifferentiated and (C) differentiated HaCaT cells nuclear extracts was assessed by the TransSignal protein/DNA combo array (Panomics, California, USA). The resulting chemiluminescence images from membranes are shown. Each dot represents the hybridization of biotinylated oligonucleotides containing consensus TF-binding sequences with immobilized complementary sequences. The intensity of signals is directly proportional to the TF binding activities present in the nuclear extracts. The full list of TFs evaluated is provided in the **S1 Table**. Dots along the right side and bottom of the membranes are controls for normalization. Images were acquired and signal densities were quantified using the ImageQuant ™ TL 8.1 software (GE Healthcare).

**S3 Fig. Confirmation of TFs overexpression with increasing input plasmids.** C-33 A cells were co-transfected with increasing amounts (25, 100 or300 ng) of p-CMV vectors expressing the respective TF, 100 ng of the recombinant vector pGL3-LCR-HPV-18-Luc, and 100 ng of pCMV-β-Gal. As a control, C-33 A cells were co-transfected with the pCMV empty vector (pCMVϕ). After total RNA extraction, the expression of each TF was evaluated by real-time PCR. Data are represented by the mean and standard deviation of a representative experiment conducted in triplicate. Three independent experiments were carried out. Statistical significance: ns, not significant, *p<0.05, **p<0.005, ***p<0.0005, ****p<0.00005.

**S1 Table. Map of transcription factors positions in the Protein/DNA ComboArray membrane (MA1215, Panomics, California, USA).**

**S2 Table. List of transcription factors with differential DNA-binding activity comparing undifferentiated and differentiated keratinocytes.**

**S3 Table. *In silico* comparison of PAX6, HMG, and NFE2 binding sites predicted positions by three different databases.**

**S4 Table. Summary of findings.**

**S5 Table. Primers used in RT-qPCR.**

**S6 Table. Primers used in chromatin immunoprecipitation (ChIP) assays.**

